# Adjustment of spurious correlations in co-expression measurements from RNA-Sequencing data

**DOI:** 10.1101/2021.03.25.436972

**Authors:** Ping-Han Hsieh, Camila Miranda Lopes-Ramos, Manuela Zucknick, Geir Kjetil Sandve, Kimberly Glass, Marieke Lydia Kuijjer

## Abstract

Gene co-expression measurements are widely used in computational biology to identify coordinated expression patterns across a group of samples, which may indicate that these genes are controlled by the same transcriptional regulatory program, or involved in common biological processes. Gene co-expression is generally estimated from RNA-Sequencing data, which are commonly normalized to remove technical variability. Here, we demonstrate that certain normalization methods, in particular quantile-based methods, can introduce false-positive associations between genes, and that this can consequently hamper downstream co-expression network analysis. Quantile-based normalization can, however, be extremely powerful. In particular when preprocessing large-scale heterogeneous data, quantile-based normalization methods such as smooth quantile normalization can be applied to remove technical variability while maintaining global differences in expression for samples with different biological attributes. We therefore developed SNAIL, a normalization method based on smooth quantile normalization specifically designed for modeling of co-expression measurements. We show that SNAIL avoids formation of false-positive associations in co-expression as well as in downstream network analyses. Using SNAIL, one can avoid arbitrary gene filtering and retain associations to genes that only express in small subgroups of samples. This highlights the method’s potential future impact on network modeling and other association-based approaches in large-scale heterogeneous data.

## 2 Introduction

Understanding the cell’s regulatory machinery can provide relevant insights into healthy tissues as well as human diseases [1, 2]. While certain experimental techniques, including chromatin immunoprecipitation sequencing (ChIP-Seq), can map interactions made by regulatory elements, such as transcription factors, it is challenging to directly observe the combined effect of multiple regulators in a systematic way. Previous studies have shown that genes undergoing similar regulatory processes tend to have coordinated expression, also called “co-expression,” across samples [3, 4, 5]. Therefore, estimates of gene co-expression are commonly used to help infer associations between genes. Gene co-expression can also be used in combination with other molecular data to improve the detection of regulatory interactions [6, 7, 8, 9, 10].

Most commonly, co-expressed genes are identified using Pearson correlation, Spearman correlation[11], and mutual information [12, 13]. Another popular approach is to first construct models that predict the expression of one gene, using that of all other genes or potential regulators, and then apply variable selection to identify the dependencies between these genes [14, 15]. Both types of approaches aim at identifying associations between genes based on their expression levels across all samples in a dataset. Therefore, as with standard gene expression analysis, it is essential to preprocess the expression data that is used as input for co-expression analysis [16].

To correct for technical variability across samples, various RNA-Sequencing (RNA-Seq) normalization methods have been developed [17, 18]. Since biological and technical variability cannot be distinguished in RNA-Seq data, algorithmic modeling is required to infer technical variability and eventually correct the read counts for this. Most normalization methods correct for technical variability using global properties (statistics that consider every sample). For instance, relative log expression (RLE) normalization, as used in DESeq computes the median ratio of gene counts relative to the geometric mean across all samples [18]. Without providing the biological group information, global shifts in gene expression caused by biological differences may be removed during the normalization process [19]. To address this issue, a quantile normalization-based method was recently developed that utilizes the information of the experimental design provided by the user to categorize samples into one or more biologically meaningful groups. Both group-specific and global properties of the expression distribution are then used to correct for technical variability. This method, called smooth quantile normalization, or *qsmooth*, yields better preservation of global shifts in expression as well as adequate control over the within-group variability [20]. While *qsmooth* was only recently developed, it has already been used in several analyses with large heterogeneous RNA-Seq datasets [1, 21, 22, 23].

Here, we show that quantile-based normalization methods, and in particular smooth quantile normalization, can introduce false-positive associations, or false-positive co-expression values, between genes. We found that this can particularly occur in datasets that have large differences in the library size across samples. To correct for these false-positives, we developed SNAIL, or **S**mooth-quantile **N**ormalization **A**daptation for **I**nference of co-expression **L**inks. SNAIL is a modified implementation of the smooth quantile normalization that uses a trimmed mean to determine the quantile distribution and applies median aggregation for genes with shared read counts (Figure 1). We analyzed RNA-Seq data from the Genotype-Tissue Expression (GTEx) Consortium [24] to showcase the problem, and data from the Mouse Encyclopedia of DNA Elements (ENCODE) [25] to validate the method. We found that SNAIL effectively removes false-positive associations between genes, without the need to select an arbitrary threshold or to exclude genes from the analysis. We anticipate that our method will benefit future co-expression and regulatory network analyses, in particular those that involve the analyses of large-scale heterogeneous RNA-Seq datasets.

**Figure 1:**
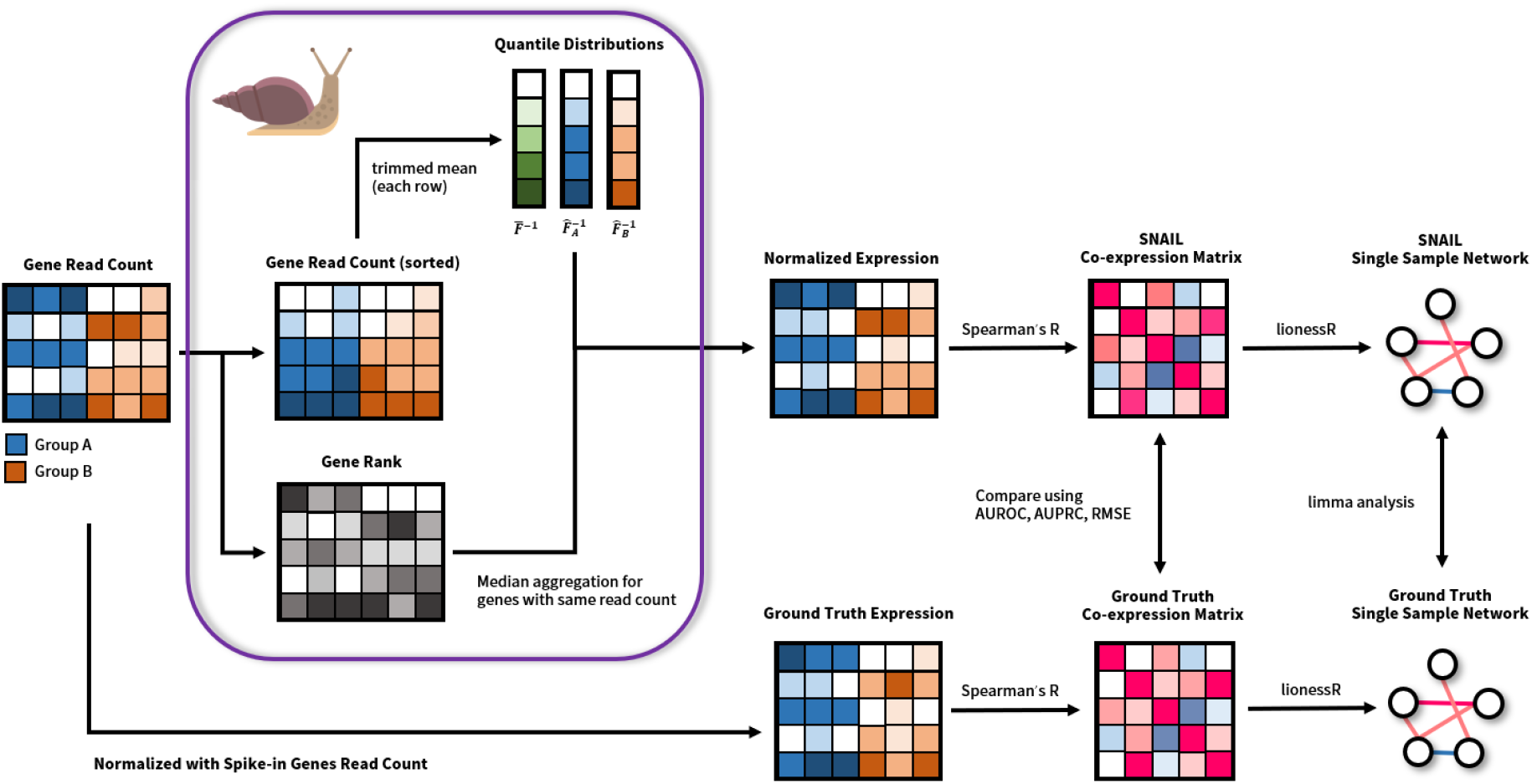
Schematic overview of SNAIL and the analyses performed in this work. SNAIL is based on *qsmooth*, but uses the trimmed mean to derive the quantile distribution for all samples as well as for every biological group of samples. In addition, SNAIL uses the median of the quantiles to normalize the expression for genes with the same read count in one sample. To evaluate SNAIL, we compared the Spearman’s correlation coefficients between genes and the edge weights in sample-specific networks with the ground truth, derived from expression normalized using spike-in genes.

## 3 Methods

### 3.1 Preparation of GTEx data

We downloaded RNA-Seq count data from the GTEx Consortium V8 release [24]. We followed the procedure conducted by Paulson *et al*., [26] to merge tissues with similar expression profiles. We selected those tissues previously reported to have the largest number of genes that deviate the most when comparing the expression in the tissue-of-interest with the median expression across all tissues—*testis, kidney* – *cortex, brain* – *other, breast – mammary tissue*, and *whole blood*) [1]. Note that the *brain – other tissue* consists of several merged brain regions, as described in Paulson *et al*., [26]. The resulting datasets consisted of 1,575 samples and 55,878 genes. Combined with the tissue information (*biosample_name*) in the meta data, we then used the Bioconductor package *qsmooth* (version 1.4.0) [20] to perform smooth quantile normalization, using tissue as the user-defined “sample group” for calculating the *group reference distributions* (see the Results section for a more detailed explanation of smooth quantile normalization). Finally, we applied our SNAIL method (see Results section) to the same dataset, again using tissue as the user-defined “sample group.”

### 3.2 Preparation of ENCODE data

For the validation datasets of our normalization approach, we downloaded bulk polyA plus RNA-seq count data consisting of twelve tissues (*embryonic facial prominence*, *forebrain*, *heart*, *hindbrain*, *intestine*, *kidney*, *limb*, *liver*, *lung*, *midbrain*, *neural tube*, *stomach*) from the Mouse ENCODE database using Bioconductor package *ENCODExplorer* (version 2.14.0, download date September 11, 2020) [27]. Among all the available experiments, we extracted those for which External RNA Controls Consortium (ERCC)-only spike-ins (accession: ENCSR884LPM) information was available. The resulting datasets consisted of 126 samples and 43,346 genes.

To establish the validation dataset, we normalized the read counts by the expression of 96 spike-in genes (Supplementary Materials 1, Section 3). Similar to the preparation of GTEx dataset (Method 3.1), we used both *qsmooth* and SNAIL to perform normalization on the original count data with corresponding tissue information of each sample (*SMTSD*) as the biological group to be used for tissue-aware normalization.

### 3.3 Evaluation of SNAIL

For both the GTEx and ENCODE datasets, we extracted the genes that were exclusively expressed in one tissue, denoted as *tissue-exclusive genes*. We define tissue-exclusitivity using the following two criteria: (1) the median ground truth expression of the gene is higher than or equal to 10 across all samples from the tissue-of-interest, and (2) the median ground truth expression is lower than or equal to 1 across samples from all other tissues. Note that we used these criteria to facilitate the visualization of the problem. We did not particularly look into the biological insights of these genes in this study.

To showcase the false-positive associations introduced by smooth quantile normalization, we compared the Spearman’s rank correlation coefficients for these tissue-exclusive genes based on qsmooth-normalized and SNAIL-normalized expression levels. Since the number of tissue-exclusive genes can show large differences across different tissues, we excluded tissue-exclusive genes of tissues with less than five or more than 1,000 tissue-exclusive genes in total from the visualization (specifically, genes exclusively expressed in testis for the GTEx dataset, and embryonic facial prominence, limb, neural tube, and forebrain for the ENCODE dataset). Note that the exclusion of these genes is not required when applying SNAIL. The number of tissue-exclusive genes for the two datasets are shown in Supplementary Materials 1 Section 4.

To evaluate the performance of SNAIL, we defined two genes to be associated when the absolute value of their Spearman’s rank correlation coefficient, based on the ground truth expression in the ENCODE dataset, was higher than or equal to a specific value. We ranged this value from 0.2 to 0.8. In addition, to evaluate whether the false-positive associations introduced by quantile normalization-based methods can propagate through downstream network analysis, we constructed sample-specific networks using Bioconductor package *lionessR* (version 1.2.0-0) [28] with Spearman’s rank correlation coefficients as the network reconstruction function. We modeled these networks using the qsmooth-normalized expression as well as using the SNAIL-normalized expression data. We then used Bioconductor package *limma* (version 3.44.1) [29] to identify significant differences of the distributions of edge weights measured across all sample-specific networks constructed from the ground truth co-expression networks. We excluded 27 tissue-exclusive genes that did not have gene symbol annotations in *biomaRt* (version 2.36.1) [30].

### 3.4 Code availability

We developed SNAIL as a Python package. The implementation of the SNAIL algorithm and all of the analyses conducted in this study can be reproduced using the Snakemake workflow management system [31] from the repository https://github.com/kuijjerlab/PySNAIL.

## 4 Results

### 4.1 Quantile-based normalization methods can introduce false-positive associations in large-scale heterogeneous datasets

In this section, we demonstrate how quantile-based normalization—and in specific smooth quantile normalization— can introduce false-positive associations between genes. To do so, we will present a case study on co-expression analysis between genes that are exclusively expressed in a specific tissue. We used RNA-Seq data from the Genotype Tissue-Expression (GTEx) project [24] and selected the tissues with high levels of tissue-specific gene expression (see Methods 3.3). We performed smooth quantile normalization to remove the technical variability presented in the dataset, while preserving the global expression differences between the different tissues. Thereafter, we extracted the tissue-exclusive genes for each tissue (see Methods 3.3) and performed co-expression analysis using Spearman’s rank correlation coefficient (*ρ*).

We would expect to only observe co-expression between genes that are expressed in the same tissue and not between genes that are exclusively expressed in different tissues. However, while we do observe high coexpression levels between tissue-exclusive genes within the same tissue, relatively high levels of co-expression are also observed between genes that are exclusively expressed in different tissues, in particular between *whole blood, lymphoblastoid cell lines* (*LCL*) and *liver* (Figure 2a). For example, when we use a threshold of *ρ*=0.3 to indicate co-expression, qsmooth normalization would introduce 3,442 false-positive associations, which is roughly 8.6% of the total number of potential associations (*n* = 40, 119).

**Figure 2:**
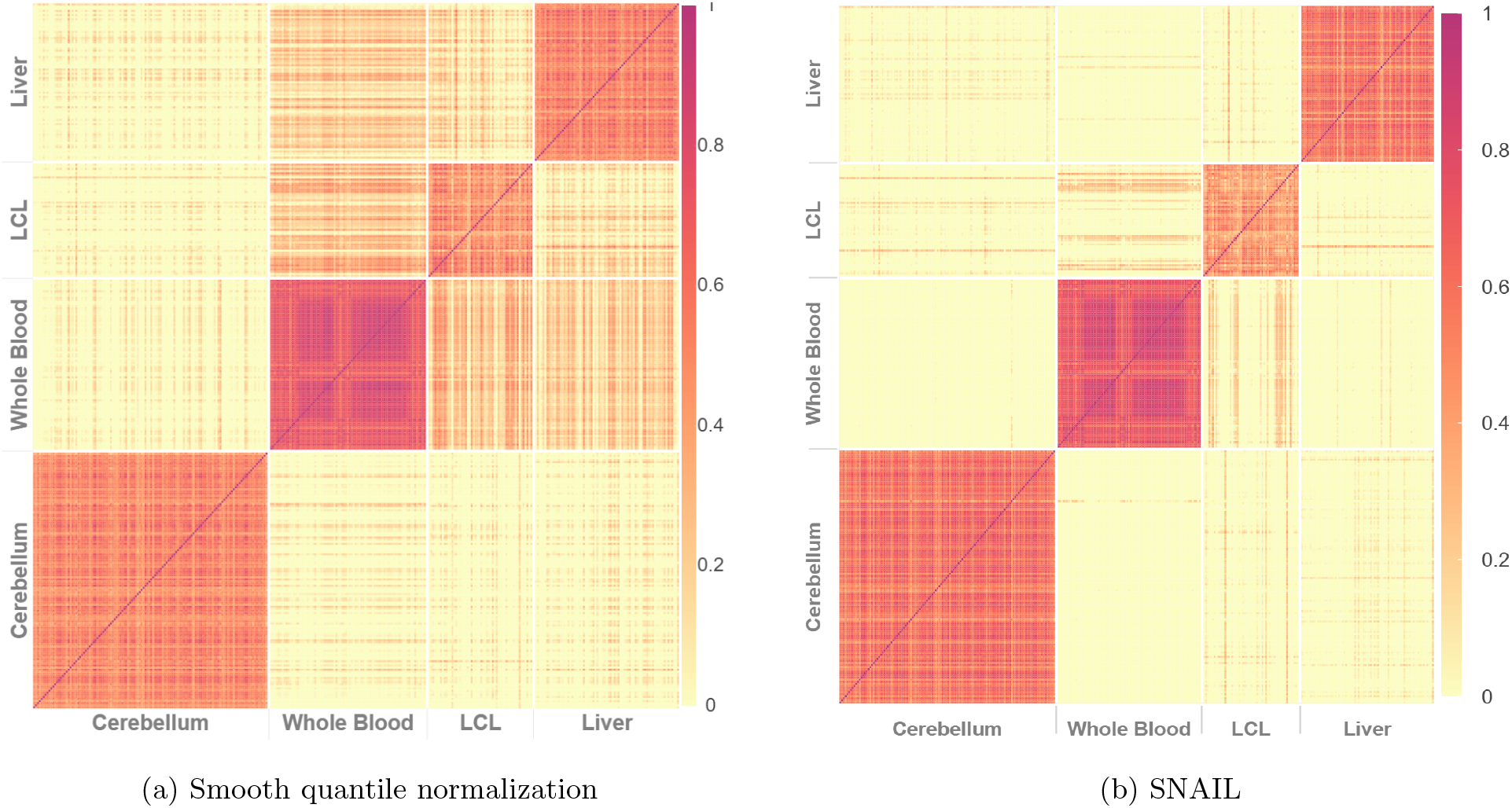
Spearman’s rank correlation coefficients between tissue-exclusive gene pairs, based on smooth quantile normalized data from GTEx. (a) False-positive associations are detected in *qsmooth*-normalized data between tissue-exclusive genes from different tissues, as can be seen in off-diagonal blocks of expression. (b) SNAIL removes most of these false-positive associations.

To understand how these false-positive associations arise, we dissected both quantile normalization (Supplementary Materials 1 Section 1, 2) and the smooth quantile normalization algorithm as it is implemented in the Bioconductor package *qsmooth* [20]. As we found the problem more prone to arise with smooth quantile normalization, we focus on this methodology in the remainder of this manuscript.

The *qsmooth* implementation of smooth quantile normalization computes the average expression level in each quantile, considering only the samples within a given user-defined group—the *group reference distribution—*as well as the average expression level in each quantile, considering all samples—the *background reference distribution*. The method then estimates the empirical reference distribution to be the weighted average of the background reference distribution and the group reference distribution, where the weight coefficient is computed based on the proportion of explained variability in the group quantile distribution. Since *qsmooth* uses the average to derive the quantile distributions, the values corresponding to small quantiles could be non-zero despite most of the values that those quantiles are based on are zeros (see also Supplementary Materials 1 Section 1, 2).

Another important detail of the implementation of quantile-based normalization is the ranking method used to process genes that have the same read count (tied counts) in each sample. The quantiles corresponding to these genes are dependent on the number of genes that have the same read count in that specific sample (Supplementary Materials 1, Section 1, 2). Therefore, even if a gene would have the exact same read count in two different samples, the corresponding quantiles can be drastically different. Especially for zero-inflated RNA-Seq data in heterogeneous datasets that have large differences between the smallest and the largest number of non-expressed genes across samples, lowly expressed genes could, for example, share the same quantile with non-expressed genes in different samples. As the normalized values are dependent on the quantiles of the expression distribution in each sample, this can introduce small technical variability across samples and this can lead to false-positive correlation coefficients between, for example, genes that only express in a subset of samples (Supplementary Materials 1 Section 1, 2).

### 4.2 Smooth quantile Normalization Adaptation for the Inference of co-expression Links

It is important to be able to take advantage of quantile-based normalization methods, and in specific smooth quantile normalization, so that one can explicitly model the biological variability and retain global expression differences in heterogeneous data. However, we also need to ensure the identified co-expression signals are reliable. We therefore developed SNAIL, or **S**mooth-quantile **N**ormalization **A**daptation for **I**nference of co-expression **L**inks (Figure 1).

SNAIL is an adaptation of smooth quantile normalization that, instead of using the average of the observed quantile distributions, uses the trimmed mean (customizable, by default, trimmed 15% largest and smallest values) to infer the heuristic reference quantile distribution (Supplementary Figure 5, step 2). In addition, when normalizing genes with the same read count, SNAIL uses median aggregation of the corresponding quantiles to substitute the original data with the normalized values. (Supplementary Figure 5, step 8). As we show below, these adaptations drastically reduce the formation of false-positive associations.

We developed SNAIL as a standalone Python package supporting multi-thread optimization. We also implemented a diagnostic function that computes the proportion of affected genes for each sample, which helps detect whether regular smooth quantile normalization would introduce false-positive associations between genes in a specific dataset (Supplementary Materials 1 Section 1).

### 4.3 SNAIL reduces false-positive associations

We applied SNAIL to normalize gene expression levels in the GTEx data and then repeated the co-expression analysis described in the previous section (see also Methods 3.3), using the same gene set and tissues. Comparing the Spearman’s rank correlation coefficients obtained in the *qsmooth*- and SNAIL-normalized data, we found that SNAIL is capable of removing false-positive associations, while modeling tissue-exclusive biological variability similarly to smooth quantile normalization (Figure 2b). With the above-mentioned threshold of *ρ*=0.3 to define co-expression, SNAIL reduces the number of such false-positive associations from 3,442 (8.6%) to 231 (0.58%).

To provide more evidence supporting the capability of SNAIL to reduce these false-positive associations, we applied the normalization method to RNA-Seq data from the Mouse ENCODE database, which includes spike-ins (Methods 3.2). We used the expression of spike-in genes to normalize the read count (Supplementary Materials 1 Section 3), creating the ground truth expression dataset. Comparing Spearman’s rank correlation coefficients obtained with *qsmooth* and SNAIL with those derived from the ground truth expression dataset for each gene pair, we observed that the root mean square error (RMSE) between the correlation coefficients decreases from 0.389 to 0.015 after applying SNAIL.

We next defined two genes to be associated if the ground truth Spearman’s rank correlation coefficients exceeded a certain threshold, which we ranged from 0.2 to 0.8. We conducted receiver-operator curve and precision-recall curve analyses and reported the area under the two curves. We found that SNAIL can reduce the false-discovery rate in co-expression analysis, regardless of the strength of the correlation signal (Figure 3). Note that when the true association is more strictly defined (correlation coefficient above 0.7), the small number of positive associations (less than 30 positive associations when the threshold is above 0.70) causes a fluctuation in the AUPRC.

**Figure 3:**
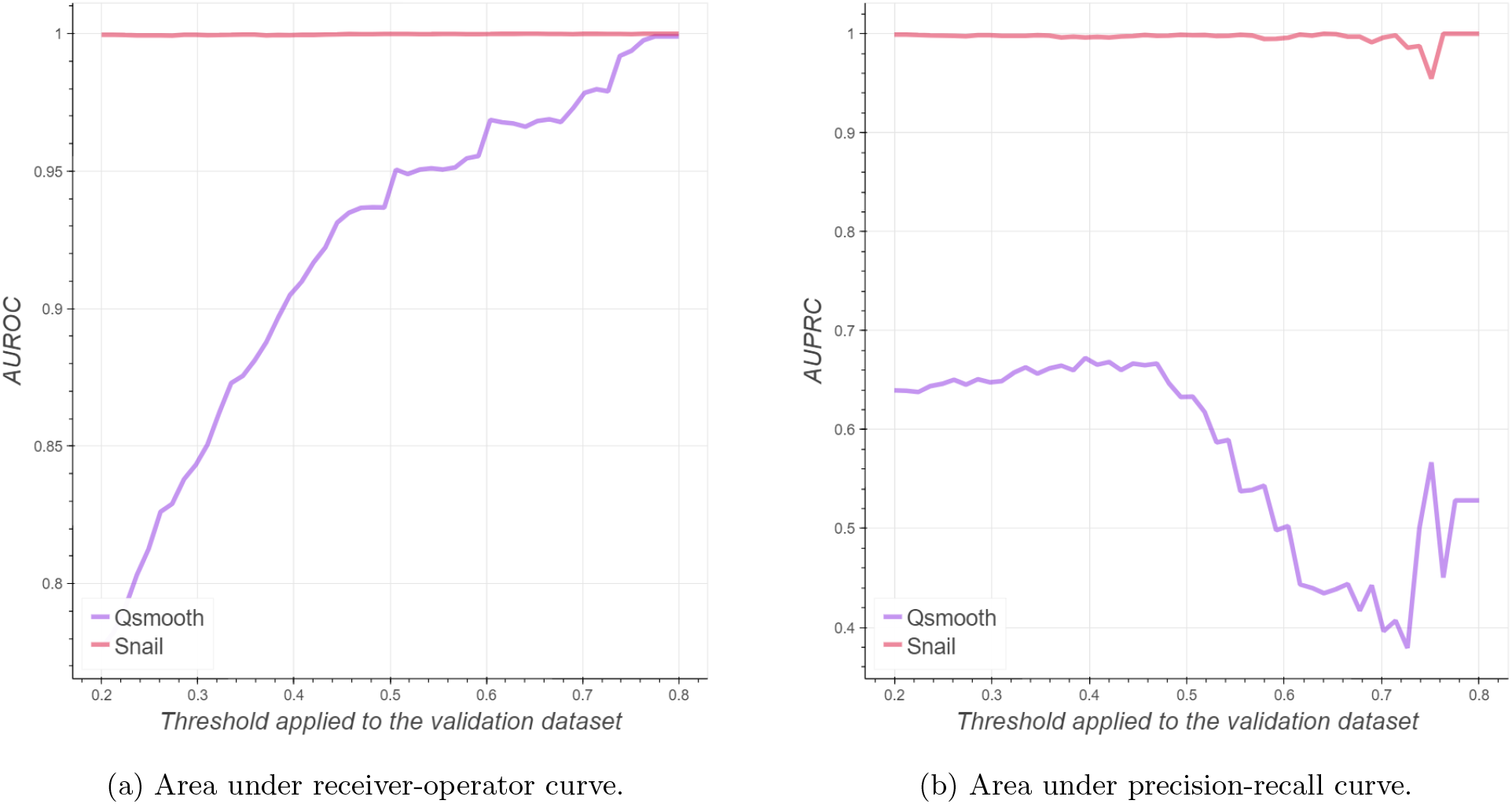
SNAIL can effectively reduce the number of false-positive associations in co-expression analysis. The x-axis denotes the threshold of absolute Spearman’s rank correlation coefficient based on ground truth expression that define true associations between genes, while the y-axis corresponds to the area under the Receiver Operator Curve (AUROC) and Precision Recall Curve (AUPRC).

In addition to these analyses, we compared SNAIL’s performance to that of other commonly used normalization methods, such as relative log expression (RLE) and, transcripts per million (TMM), (Supplementary Materials 1 Section 5). Compared to *qsmooth*, SNAIL effectively removes false-positive associations, while reaching a similar performance in detecting correct associations between genes as RLE and TMM (Supplementary Figure 6). Note however, that the performance of these methods can not be directly comparable, as SNAIL and smooth quantile normalization explicitly model the global differences across different biological groups and show better control for the within-group variability (Supplementary Figure 7). The comparisons we made here aim to showcase the limitation of the original implementation of smooth quantile normalization when normalizing data to be used in correlation analyses.

### 4.4 SNAIL reduces false-positives in downstream network analysis

We next wanted to evaluate whether the false-positive associations introduced by quantile-normalized methods also affect downstream network analysis. To explore this, we used the LIONESS algorithm [32] to infer sample-specific networks from co-expression measurements. LIONESS is based on the assumption that edges estimated in an “aggregate” network model are a linear combination of edges specific to each of the input samples. This allows for the estimation of individual sample edge weights using a linear equation, these can then be used for sample-specific network analysis, as done previously [33, 34, 35].

The LIONESS algorithm constructs a network for each sample, where the nodes denote the genes and edge weights denote the likelihood of two genes being associated. We compared the distribution of edge weights across all sample-specific networks constructed on ground truth co-expression to the distribution of edge weights from the co-expression networks constructed on (a) *qsmooth*- and on (b) SNAIL-normalized expression, using a t-test for each gene pair independently. Figure 4a shows that the false-positive associations propagate through downstream network analysis, creating 199 false-positive edges from a total of 1,484 potential gene pairs between genes exclusively expressed in different tissues. In SNAIL-normalized data, no edges significantly differ from the network built on the ground truth expression (FDR adjusted P-value ≤ 0.001, Figure 4b). This analysis shows that SNAIL can effectively remove false-positive associations between genes, and can thus improve the reliability of downstream network analysis.

**Figure 4:**
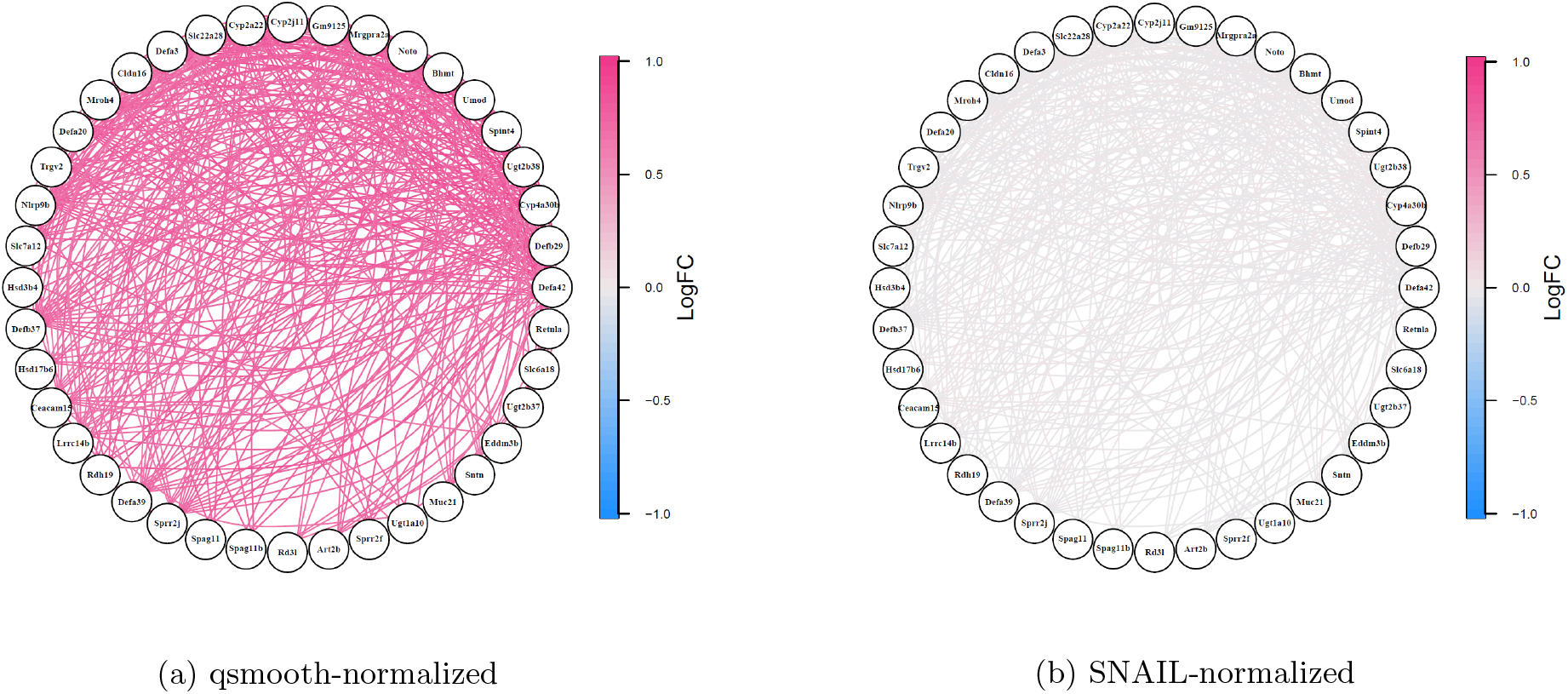
Log-transformed fold changes on the mean value (computed across sample-specific networks) of the edge weights comparing the network constructed on ground truth expression with the networks constructed on (a) the smooth quantile-normalized expression and on (b) the SNAIL-normalized expression levels. All edges with a significant difference (FDR-adjusted P-value ≤ 0.001) between the ground truth and either of the normalization methods are shown. The edge weights denote the likelihood of two genes being associated in each sample. The log-transformed fold change between the edge weight is color-coded, with magenta indicating that the edge weights between the two genes are over-estimated in the sample specific networks constructed using the dataset with evaluated normalization method compared to the ground truth, and blue indicating under-estimation of the edge weights.

## 5 Discussion

Here, we showed that the use of quantile-based normalization approaches—and in specific smooth quantile normalization—to RNA-Seq data can introduce false-positive associations between genes, and that this can propagate to and affect downstream network analyses. We found that false-positive associations particularly arise when there is a large difference between the smallest and the largest number of non-expressed genes across the samples in the dataset. This can for example occur when dealing with RNA-Seq datasets collected from large-scale projects that include heterogeneous data, such as data from The Cancer Genome Atlas (TCGA) [36], ENCODE, and GTEx.

A frequently applied strategy that attempts to remove potential false-positive associations is filtering out genes with low read counts across a certain number of samples. However, thresholds used for filtering are often chosen arbitrarily, and can remove genes that are specifically expressed in a subset of samples, such as the tissue-exclusive genes that we described in our example of network analysis in data from GTEx. Therefore, arbitrary filtering is not ideal if one aims to compare gene associations or networks derived from different subgroups of samples. Moreover, it would be ideal to include all genes in large-scale network analysis, as certain network reconstruction algorithms make use of the entire distribution of gene expression and thus filtering out genes may remove some signal from the input dataset [6].

To address this issue, we developed SNAIL, an adaptation of smooth quantile normalization, which retains global expression differences in heterogeneous datasets through the use of both group-specific and global properties. By using the trimmed mean to infer the reference quantile distribution as well as median aggregation for genes with the same read count, SNAIL avoids the formation of false-positive associations. Importantly, as the method does not require gene filtering, it allows for direct comparison of networks modeled on heterogeneous datasets.

While we specifically focused our examples on modeling co-expression across different tissues in this work, false-positive associations can also arise when comparing other biological conditions that show large differences in expression profiles under certain experimental settings, such as when comparing networks for males and females [34]. We also envision that other methods that are based on correlations, such as eQTL studies, could potentially include quantile-based normalization-introduced false-positives, and could benefit from normalization with SNAIL. In general, we would like to raise awareness of implementing tools designed for gene expression data in existing correlation-based approaches or pipelines. Most of the published evaluations of normalization methods are based on comparing differences between ground truth and normalized expression levels. However, the impact of normalization on correlation-based measures is often neglected.

Heterogeneous datasets with increasing numbers of samples and conditions will likely be published in the near future, and new methods for combining data from different studies [37] will result in the emergence of even larger and more heterogeneous datasets. As these datasets will become available for analysis, SNAIL will be an important tool that will allow for more precise analyses of large-scale data with network-based approaches.

## 7 Acknowledgments

The authors would like to thank the Kuijjer and Mathelier groups for helpful discussions and Elisa Bjørgø and Ingrid Kjelsvik for administrative support.

## 8 Author contributions

Conceptualization, MLK, CML, KG; Methodology, PHH, MLK; Software, PHH; Formal Analysis, PHH; Writing — Original Draft, PHH, CML, MLK; Writing — Review Editing, CML, GKS, KG, MKK, MZ; Supervision, MKK; Funding Acquisition, MKK.

