## Supplementary Materials 1 for "Adjustment of spurious correlations in co-expression measurements from RNA-Sequencing data"

### Adjustment of spurious correlations in co-expression measurements from RNA-Seq: Supplementary Material 1

June 23, 2022

#### 1 Dissection of quantile normalization

Quantile normalization starts with deriving the empirical reference distribution $\bar{F}^{-1}(u)$ using the average of quantile distribution across all samples $F_j^{-1}(u)$:

$$\bar{F}_j^{-1}(u) = \frac{1}{n}\sum_j F_j^{-1}(u)$$

, where $n$ is the number of samples and $j$ is the sample index. To normalize the read count for each gene, quantile normalization replaces the observed data distribution $G$ of each sample with the empirical reference distribution by substituting the read count of the original dataset with the values that share the same order in the empirical reference distribution:

$$X^{(norm)} = \bar{F}^{-1}(G(X)) \ .$$

Here, we describe how quantile normalization can give rise to false-positive associations between genes. Given a read count data $\mathbf{X}$ and corresponding ranks for each value in the read count matrix $\mathbf{R}$ (with average ranking):

$$\mathbf{X} = [x_{ij}] \in \mathbb{N}_0^{p\times n},\ \mathbf{R} = [r_{ij}] \in \mathbb{N}_0^{p\times n}$$

, where $p$ is the number of genes, $n$ is the number of samples, $i$ is the gene index, and $j$ is the sample index, then, to derive the empirical reference distribution, quantile normalization starts from sorting the read counts in $\mathbf{X}$ for each sample (column), with the sorted read count denoted as $\mathbf{S} = [s_{uj}] \in \mathbb{N}_0^{p\times n}$ (Supplementary Figure 1, step 1), where $u$ is the rank index. The procedure continues by computing the average for each row of $\mathbf{S}$ (Supplementary Figure 1, step 2 and 3):

$$RankAverage(u) = \frac{1}{n}\sum_{j=1}^{n} s_{u,j}$$

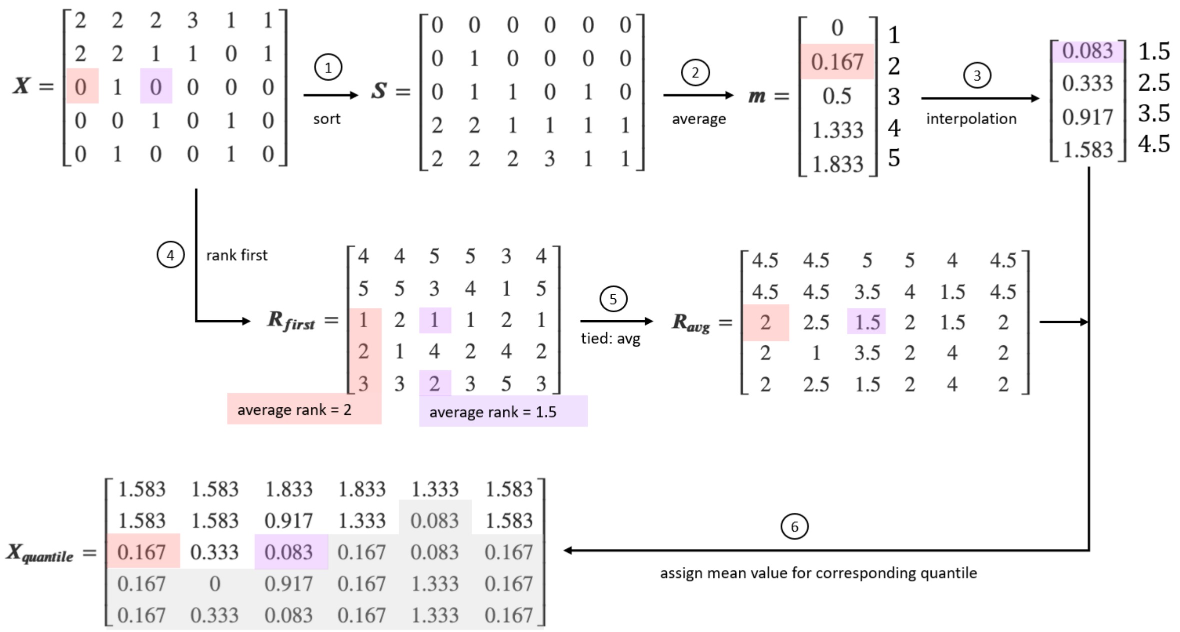


Supplementary Figure 1: Quantile normalization. The method starts from sorting each column in the count matrix (Step 1), then takes the average for each row and interpolates the values between each entry to derive the empirical reference (Step 2 and 3). To correct the count matrix, it uses the average ranking method to rank each column in that matrix (Step 4 and 5) and replaces the original values with the empirical reference using the rank as the index (Step 6). The highlighted colors are two examples showing how quantile normalization add technical variability to non-expressed genes. The orange color highlights the process when the rank is an integer, while the purple highlights the process when the rank is not an integer (and therefore requires the use of the interpolated reference to correct the count).

Finally, the procedure substitutes the read counts with the mean value of the corresponding quantile (Supplementary Figure 1, step 6):

$$x_{i,j}^{(norm)} = f(r_{i,j})$$

, where $f$ is a transformation that maps a rank to its mean value across a group of samples:

$$f(r) = \begin{cases} RankAverage(r), & r \in \mathbb{N}_0 \\ 0.5 \times (RankAverage(r-0.5) + RankAverage(r+0.5)), & \text{otherwise.} \end{cases}$$

There are two important details that can lead to false positive associations between genes. First, if we apply quantile normalization to zero-inflated gene expression data, when computing the $RankAverage(u)$ value corresponds for a small rank $u$, as long as there exists one non-zero entry across all columns (samples), the resulting value used to represent rank $u$ will be a non-zero value, depending on the sum of all non-zero entries (e.g. the second entry in $\mathbf{m}$ in Supplementary Figure 1 step 2).

Another important detail is the substitution of the normalized expression for genes with the same read count in each sample. Suppose that the $i^{th}$ to $(i+k)^{th}$ entries in the $j^{th}$ column vector (sample) of $\mathbf{S}$ have the same read count. Then, the ranks recorded in $\mathbf{R}$ corresponding to these entries will be $\mathbf{R}_{i:i+1,j} = (2i+k)/2$ (Supplementary Figure 1 step 5). This is often referred to as the *average* ranking method when dealing with tied values. If the rank is not an integer, the normalized expression value will be $0.5 \times (RankAverage(r - 0.5) + RankAverage(r + 0.5))$. For low reads in the count matrix, it is common to have the same number of counts with many other entries due to the zero-inflated nature of RNA-Seq data. The ranks corresponding to these values will be highly dependent on the number of entries with the same values ($k$) in that particular sample $j$.

Consider we have a bulk RNA-Seq dataset with 1,000 samples that includes two genes that exclusively express in different subsets of samples, say samples 600-800 and samples 800-1,000, respectively. The quantile normalized values for these two genes will be exactly the same in each sample for samples 1-600, since neither of these genes are expressed in these samples. However, since the normalized values depend on the rank of each sample, the normalized gene expression across samples can be different. When computing the Spearman's rank correlation coefficient between these two genes, the rank of the samples will be different for samples 1-800, despite that the original read counts are the same. Moreover, the rank of the samples for the two genes will be exactly the same, and this would increase the Spearman's rank correlation coefficient, leading to false-positive associations between these two genes (Supplementary Figure 2).

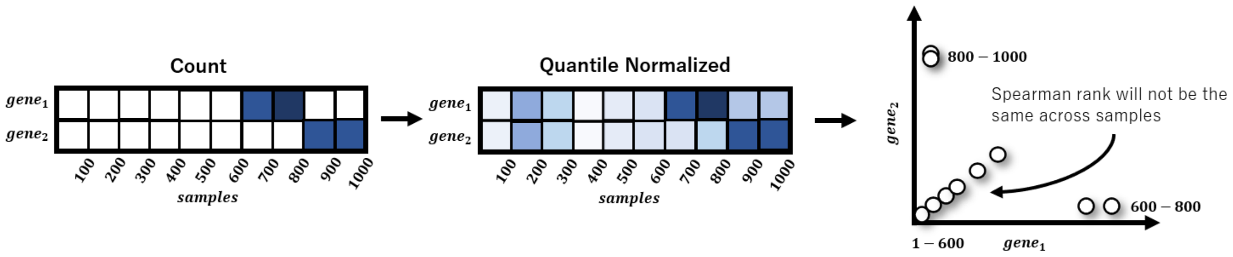


Supplementary Figure 2: Schematic example of false positive associations introduced by quantile-based normalization. Here, we visualize two genes that express in different sets of samples. We therefore do not expect these two genes to be co-expressed. However, as quantile-based normalization will add small variability to the count data, especially for entries with low (zero) read counts, this will affect the Spearman ranks of the genes, and thereby lead to the formation of false-positive associations between these two genes, as can be appreciated from the figure on the right.

The above mentioned problem is therefore more likely to happen in large-scale heterogeneous datasets where the library size is not uniformly distributed—as, for instance, in the GTEx and ENCODE datasets that we analyzed in this study (Supplementary Figure 4)—as the result of $RankAverage(u)$ will be sensitive to outliers for low quantiles. Eventually, this leads to false positive associations between genes that exclusively express in a small proportion of samples. Note that this can also happen for other lowly-expressed genes, depending on how many genes share the same read count in one sample.

Knowing this issue, we can develop a diagnostic function to detect whether false-positive associations will form between genes. If we compute the number of non-expressed genes for each sample, denoted as a vector $z$:

$$\mathbf{z} = \begin{bmatrix} z_1 & z_2 & \cdots & z_n \end{bmatrix}^T .$$

The rank $r$ for these non-expressed genes $ne$ in each sample will then be

$$\mathbf{r}^{(ne)} = \begin{bmatrix} (z_1 + 1)/2 & (z_2 + 1)/2 & (z_n + 1)/2 \end{bmatrix}^T .$$

When $z_j > 2 \times \min(\mathbf{z}) - 1$, the normalized expression value for all the non-expressed genes of $sample_j$ will be a non-zero value, and this value will again depend on the number of non-expressed genes in the sample. While we specifically focused our examples here on genes with zero read counts. The false-positive associations can also arise in lowly-expressed genes (Supplementary Figure 3).

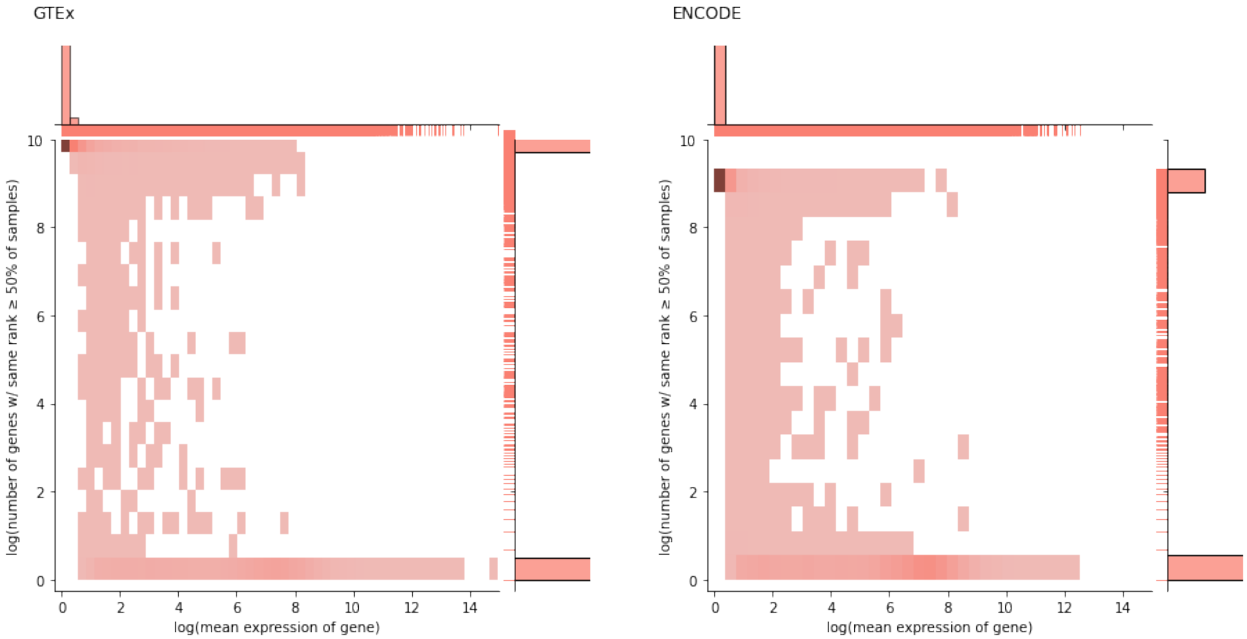


Supplementary Figure 3: The number of genes with shared rank. The X-axis denotes the log-transformed mean expression across sample for each gene. The Y-axis denotes the log-transformed number of genes with the same rank across more than 50% of the samples.

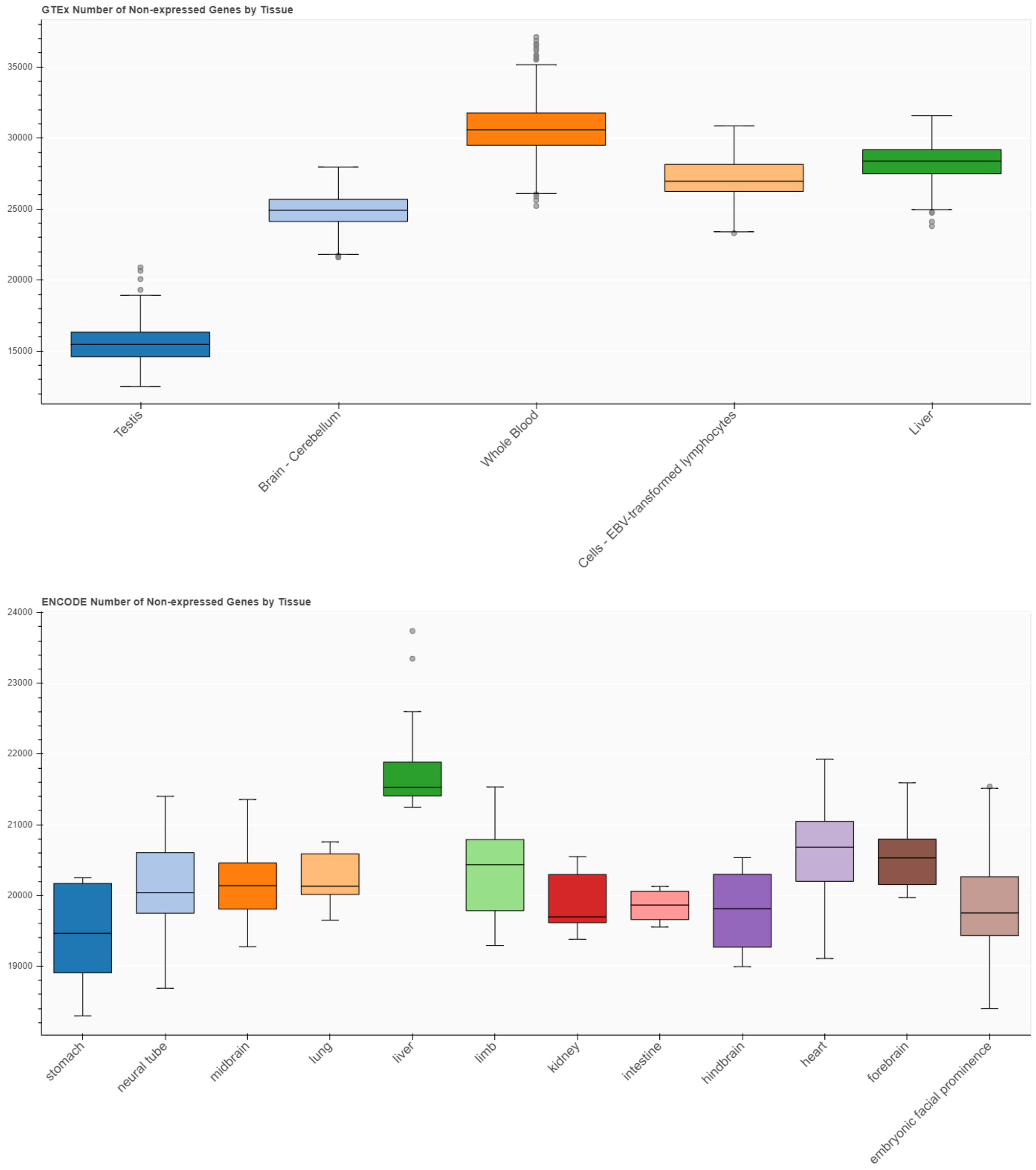


Supplementary Figure 4: Number of non-expressed genes in each sample.

### 2 Dissection of the smooth quantile normalization algorithm

Smooth quantile normalization assumes the observed quantile distribution $F_j^{-1}(u)$ to be the affine transformation of known covariates, such as the variability explained by biological groups $Z_j(u)$:

$$F_j^{-1}(u) = Z_j(u)\beta_j + \varepsilon_j(u)$$

, where $u$ is the index for the quantiles, $j$ is the index for the samples, and $\varepsilon_j(u) \sim N(0, \sigma^2)$. In addition, smooth quantile normalization assumes a similar affine transformation exists within each biological group:

$$\hat{F}_j(u)^{-1} = Z_j(u)\hat{\beta}_j + \varepsilon_j(u)$$

, where $\hat{F}_j^{-1}(u)s$ is the group reference distribution. By fitting a regression model between the observed quantile distribution $F_j^{-1}(u)$ and the group reference distribution $\hat{F}_j^{-1}(u)$, smooth quantile normalization derives the estimated heuristic reference distribution as follows:

$$F_j^{qsmooth}(u) = w_u \bar{F}_j^{-1}(u) + (1 - w_u)\hat{F}_j^{-1}(u)$$

,where

$$w_u = median(1 - \frac{SSB_u}{SST_u})$$

and $\bar{F}_j^{-1} = \frac{1}{N}\sum_j F_{uj}^{-1}$ is the average of the observed quantile distribution. $SSB$ denotes the explained sum of squares and $SST$ denotes the total sum of squares:

$$\sum_{j=1}^{n}(F_j^{-1}(u) - \bar{F}^{-1}(u))^2 = \sum_{j=1}^{n}(F_j^{-1}(u) - \hat{F}_j^{-1}(u))^2 + \sum_{j=1}^{n}(\hat{F}_j^{-1}(u) - \bar{F}^{-1}(u))^2 \ .$$

Lastly, smooth quantile normalization replaces the observed data distribution $G$ of each sample with the heuristic reference distribution by substituting the read count of the original dataset with the values that share the same order in the heuristic reference distribution. Smooth quantile normalization compares the biological variability within each given group ($\hat{F}^{-1}$) and the background ($\bar{F}^{-1}$) and use this information to compute the weighted average for means of each quantile in each group (Supplementary Figure 5 step 5). This is particularly useful when we want to preserve global shifts in the distribution of expression across different biological groups of samples, such as different tissues.

Here, we specifically discuss smooth quantile normalization implemented in Bioconductor package *qsmooth*. Given expression data from samples of $K$ different groups, then smooth quantile normalization computes the average value of each quantile considering only those samples belonging to a given group, as well as the average value of the quantile considering all samples:

$$GroupRankAverage(i, k) = (\frac{1}{n_k}\sum_{j \in N^{(k)}} s_{i,j})$$

, where $N^{(k)}$ denotes the set of sample indices in group $k$ ($N^{(0)}$ denotes indices of all samples), and $n_k$

denotes the number of samples in group $k$ (Supplementary Figure 5, step 2). The procedure continues by computing the explained sum of squares $SSB$ and the total sum of squares $SST$:

$$\mathrm{SSB}_i = \sum_{k=1}^{K} (GroupRankAverage(i,0) - GroupRankAverage(i,k))^2$$

$$\mathrm{SST}_i^{(g)} = \sum_{j=1}^{n} (GroupRankAverage(i,g) - GroupRankAverage(i,0))^2$$

, where $g$ denotes the group index for sample. The weight can then be calculated using $SSB$ and $SST$ (Supplementary Figure 5, step 6). Finally, the normalized expression value $m$ assigned back is the weighted average of the means derived from the sample group and from the background of the corresponding rank based on the weight coefficient $w$ (Supplementary Figure 5. step 7):

$$w_i = \mathrm{smooth}(1 - \frac{SSB_i}{SST_i})$$

$$m'_i = w_i GroupRankMean(i,0) + (1 - w_i) GroupRankAverage(i,j)$$

, where $j$ is the group index of the sample. Another difference between the implementation of smooth quantile normalization and quantile normalization is the way these methods process genes with the same read count. Given the $i^{th}$ to $(i+l)^{th}$ entries of the sorted expression matrix from one particular sample $\mathbf{S}_j$ have the same read count, then the ranks recorded in $\mathbf{R}$ of the genes corresponding to these entries will be randomly selected from $\{i, i+1, ..., i+l\}$ without replacement. (Supplementary Figure 5, step 3). This is often referred to as the *random* ranking method when dealing with tied values. The normalized expression for these genes will be (Supplementary Figure 5, step 8):

$$\mathbf{X}_{i:i+l,j}^{(norm)} = \frac{1}{l} \sum_{a=i}^{i+l} m'_a \ .$$

Similar to quantile normalization, smooth quantile normalization suffers from the problem of tied values presented in the data and can introduce non-zero values when computing the $GroupRankAverage(i,j)$. This also leads to false positive associations when using Spearman's rank correlation coefficient.

### 3 Validation dataset

Using the read counts of 96 spike-ins genes $\mathbf{X}^{(spikein)}$, we established the validation dataset with the following equation:

$$\mathbf{x}_j^{(val)} = \frac{\mathbf{x}_j^{(count)}}{s_j}, \quad s_j = \frac{n \sum_{i=1}^{s} x_{ij}^{(spikein)}}{\sum_{k=1}^{n} \sum_{i=1}^{s} x_{ik}^{(spikein)}}$$

, where $n = 126$ is the number of samples, $s = 96$ is the number of spike-in genes, and $\mathbf{x}_j$ denotes the column $j$ vector in the matrix $\mathbf{X}$.

#### 4 Number of tissue exclusive genes

| ENCODE | | GTEx | |
|---|---|---|---|
| Kidney | 27 | Testis* | 3992 |
| Intestine | 13 | Brain - Cerebellumn | 122 |
| Lung | 12 | Whole Blood | 81 |
| Heart | 11 | Cells - EBV-transformed lymphocytes | 54 |
| Liver | 11 | Liver | 75 |
| Stomach | 9 | | |
| Embryonic Facial Prominence* | 3 | | |
| Limb* | 1 | | |
| Neural Tube* | 1 | | |
| Forebrain* | 1 | | |

* Excluded from analysis for visualization purposes.

#### 5 Comparison with other normalization methods

To further compare the performance of SNAIL with other normalization methods, we used the same validation dataset from ENCODE and repeated the analysis as described in the Method 3.3 of the main manuscript. In addition to *qsmooth* and SNAIL, we applied relative log expression (RLE) and trimmed mean of M values (TMM) normalization using R package *NormExpression* (version 0.1.0) with default settings. Here, in addition to using tissue-exclusive genes, we performed the co-expression analysis on all genes.

We next defined two genes to be associated if the ground truth Spearman's rank correlation coefficients exceeded a certain threshold, which we ranged from 0.2 to 0.8. We conducted receiver-operator curve (Supplementary Figure 6 left panel) and precision-recall curve analyses (Supplementary Figure 6 right panel) and reported the area under the curves. The results show that SNAIL can achieve similar performance compared to other normalization methods.

It is important to note, however, that the performance of these different methods can not be directly compared, as SNAIL and *qsmooth* explicitly model the global differences across different biological groups and show better control for the within-group variability (Supplementary Figure 7).

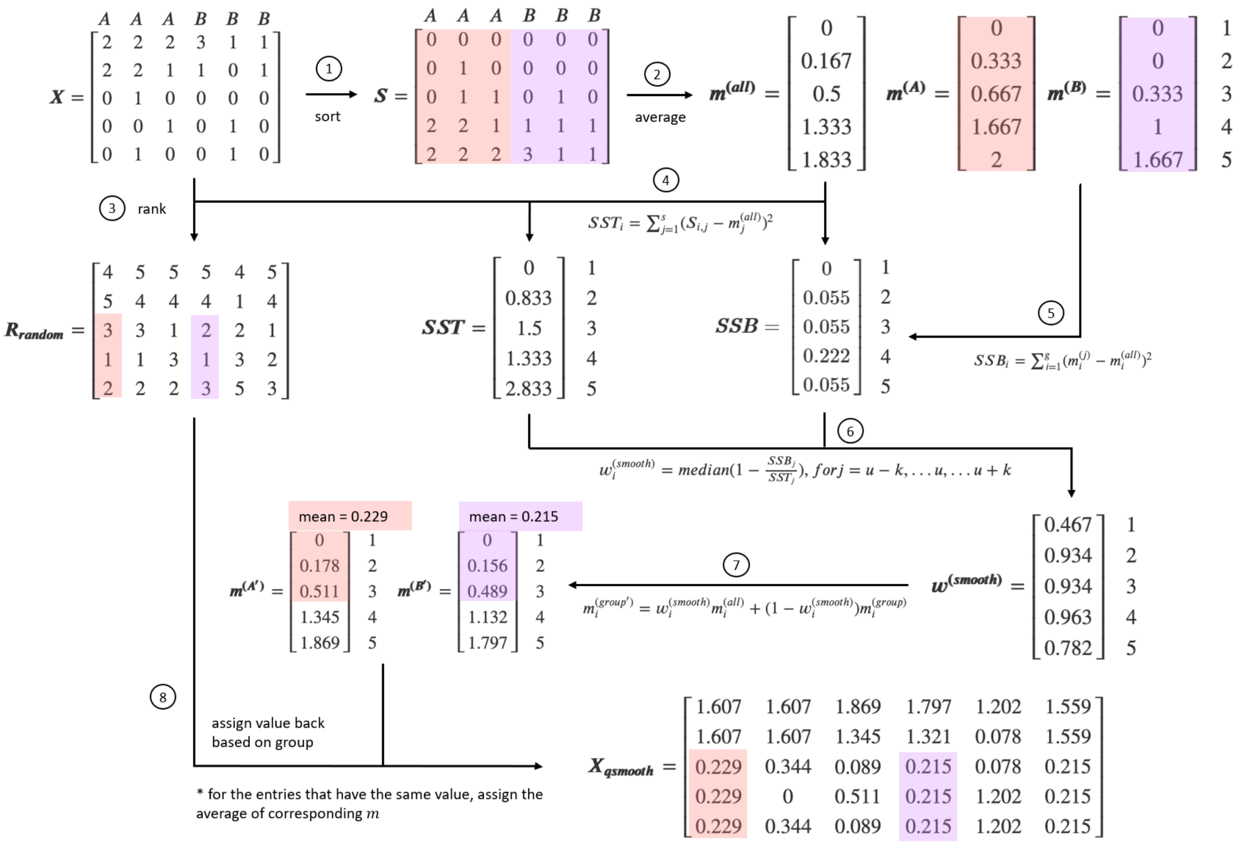


Supplementary Figure 5: Smooth Quantile normalization. $A$ and $B$ represent the group information provided by the user (for example, two different tissues). Smooth quantile normalization starts from computing the empirical reference considering all samples, as well as each group separately (Step 1 and 2). Thereafter, it uses the total sum of squares and explained sum of squares to derive the weight coefficient (Step 4 - 6). To correct the count matrix, it uses the average ranking method to rank each column in the count matrix (Step 3) and replaces the original values with the weighted sum of the empirical reference considering all samples and the group-specific samples using the rank as the index (Step 7 - 8). The colors highlight the correction process for entries in two different groups.

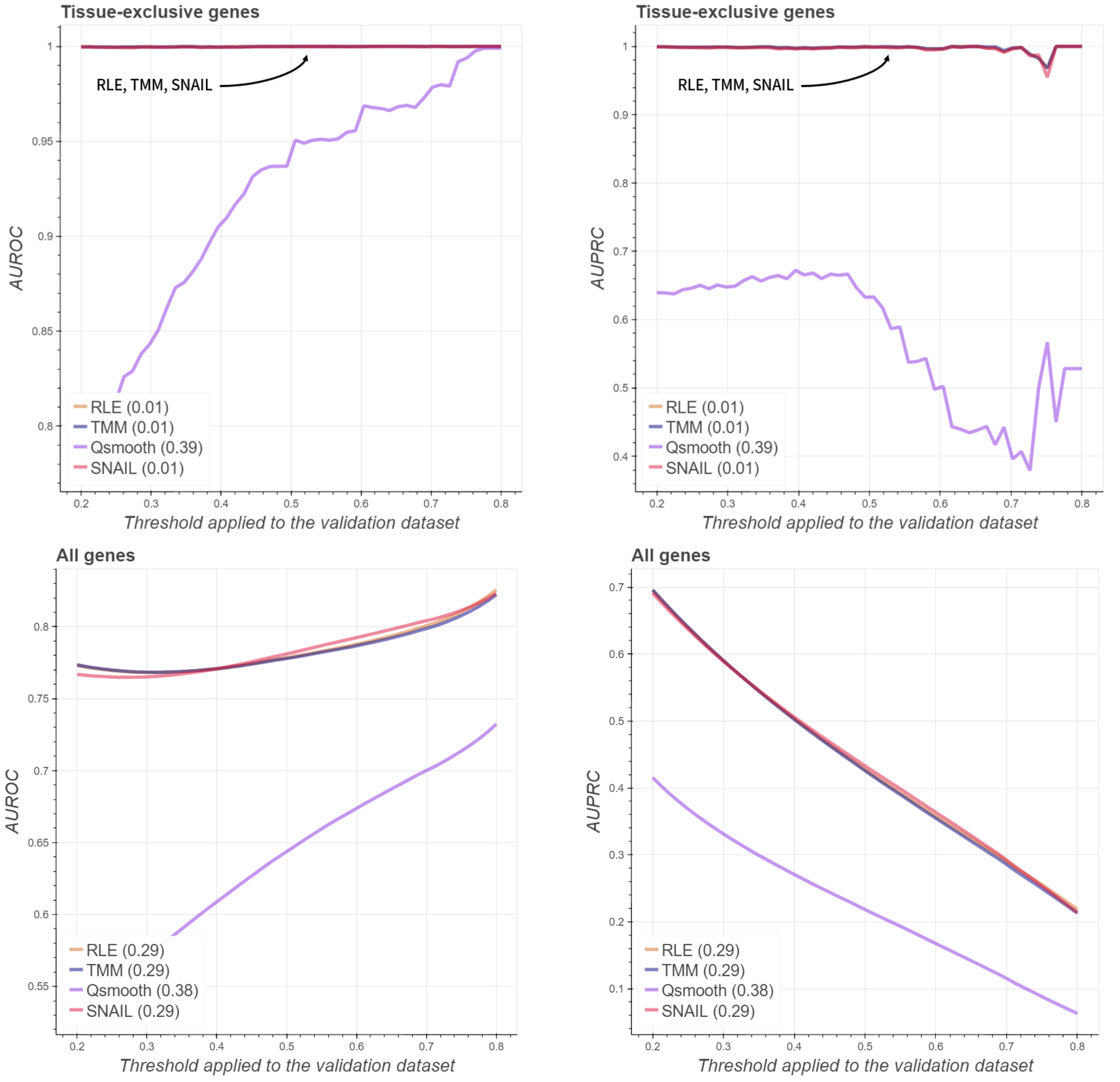


Supplementary Figure 6: Area under receiver-operator curve compared with other normalization methods. The x-axis denotes the threshold of absolute Spearman's rank correlation coefficient used to define true associations between genes in the validation dataset, while the y-axis corresponds to the area under curve. The numbers next to the normalization method in the figure legend represent the root mean squared error.

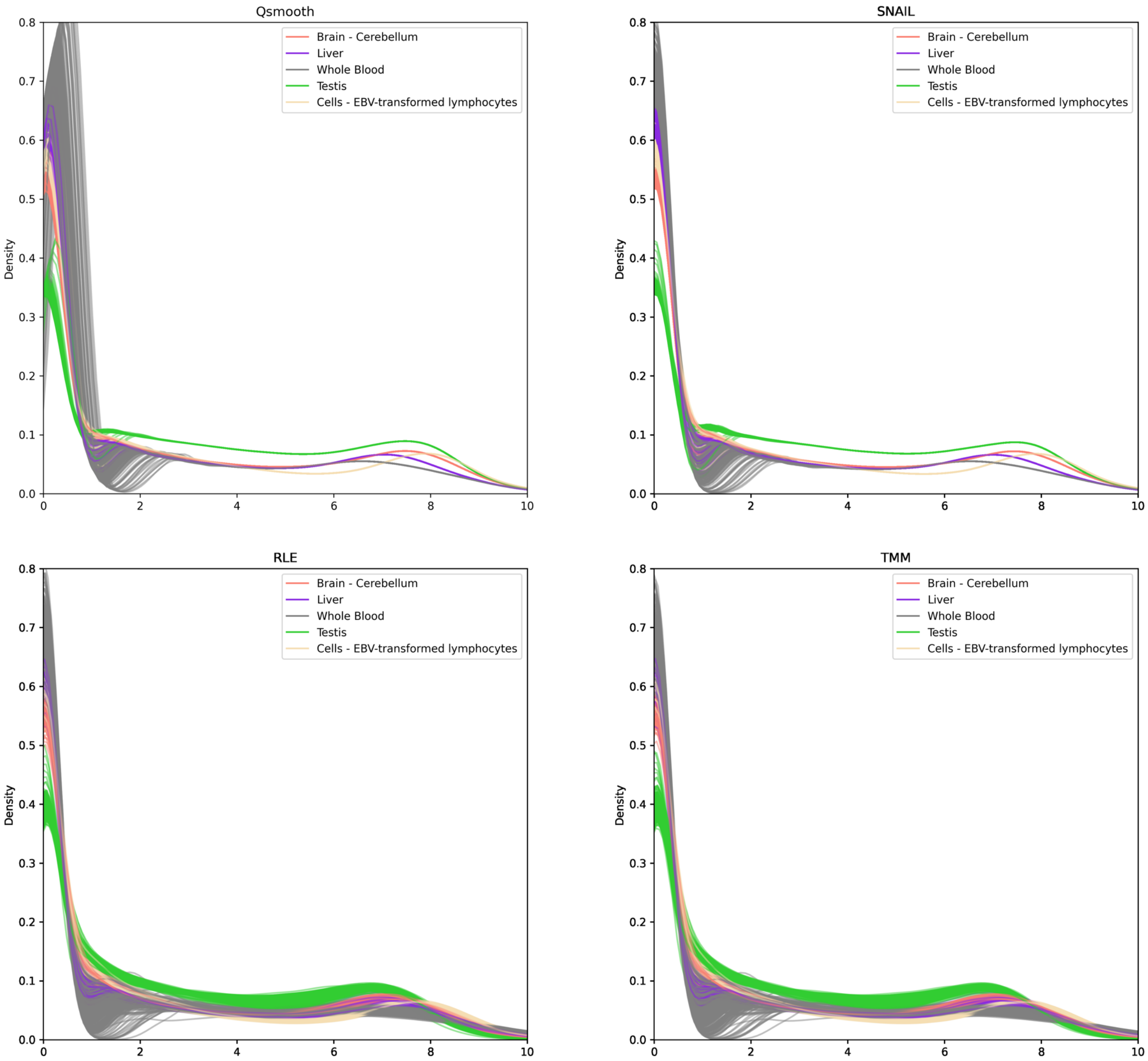


Supplementary Figure 7: The distribution of gene expression across all samples for different normalization methods on GTEx dataset. SNAIL and qsmooth show better control for the within-group variability.
